# Full inter-hemispheric integration sustained by a fraction of posterior callosal fibers

**DOI:** 10.1101/2025.02.14.638327

**Authors:** Tyler Santander, Selin Bekir, Theresa Paul, Jessica M. Simonson, Valerie M. Wiemer, Henri Etel Skinner, Johanna L. Hopf, Anna Rada, Friedrich G. Woermann, Thilo Kalbhenn, Barry Giesbrecht, Christian G. Bien, Olaf Sporns, Michael S. Gazzaniga, Lukas J. Volz, Michael B. Miller

## Abstract

The dynamic integration of the lateralized and specialized capacities of the cerebral hemispheres constitutes a hallmark feature of human brain function. This inter-hemispheric exchange of information critically depends upon the corpus callosum. Classical descriptions of callosal organization outline a topographic gradient, such that specific fibers integrate distinct aspects of brain function. Here we present a challenge to this conventional model. Using neuroimaging data obtained from a new cohort of adult corpus callosotomy patients, we leverage modern network neuroscience techniques to show—for the first time—that full inter-hemispheric integration can be achieved via a small proportion (∼1 cm) of intact posterior callosal fibers. Only complete callosotomy patients demonstrated the expected dissolution of typical inter-hemispheric network architectures, aligning with disconnection syndromes long-thought to reflect diminished information propagation and communication across the brain. These findings motivate a novel mechanistic understanding of synchronized inter-hemispheric neural activity for large-scale human brain function and behavior.

## Main

In recent decades, network neuroscience has emerged as a powerful analysis framework for modeling functional interactions between remote neural systems. A foundational assumption of this approach is that anatomical links between brain regions constrain neuronal dynamics and support the emergence of large-scale functional networks, which in turn enable the efficient integration of perceptual, motor, and higher-order cognitive information necessary for human behavior^1,2,3,4,5,6^. The corpus callosum (CC) is thought to be the central conduit through which specialized processing nodes in each cerebral hemisphere coordinate and synchronize function, with 200-250 million axons enabling direct, long-range communication across the brain. Classical topographic models of CC projections have revealed a well-defined anterior-to-posterior anatomical gradient, corresponding to the spatial organization of canonical brain networks supporting various sensorimotor and cognitive functions^7,8^. One consequence of this view is the prediction that damage to the CC should result in drastic functional disconnections between hemispheres—with disruptions following the localization of lesioned fibers along the aforementioned anatomical gradient.

Split-brain patients, in whom the CC has been severed in an effort to treat medically-intractable epilepsy, offer a powerful means of testing this core conceit of structure-function correspondence. However, neural data on these patients remain scarce—callosotomy was largely discontinued around the turn of the 21^st^ century, coinciding with the rise of network neuroscience. Extant data largely stem from anesthetized pediatric patients^9,10,11,12^. Taken together, these studies support the hypothesis that full versus partial callosotomy confer differential effects with respect to the location and extent of functional disconnections. At the same time, however, they have provided contradictory findings regarding recovery of function post-operatively^9,11,12^, and various other methodological factors obfuscate more straightforward interpretations. Thus, to date, it remains unknown how severing the largest fiber bundle of the human brain affects the topological organization and intrinsic dynamics of large-scale functional networks.

To address this fundamental question, we present the first investigation characterizing the impacts of full and partial callosotomy on the adult human brain using multiple network neuroscience methods. We hypothesized that full callosotomy would wholly disrupt functional synchrony across the cerebral hemispheres, while the spatial distribution of functional disconnections following partial callosotomy would unveil the anatomical gradient of the CC. As predicted, we found that complete sectioning of the CC broadly obliterated both typical inter-hemispheric network topologies and dynamic patterns of inter-hemispheric temporal synchrony. More strikingly, however, we also found that a small proportion of intact posterior CC fibers (a case of partial callosotomy) was sufficient to support widespread inter-hemispheric coupling. These findings challenge our conventional understanding of the CC’s functional topography and sheds new light on the critical mechanisms for inter-hemispheric information integration.

## Patient descriptions

Six adult callosotomy patients (5 men, 1 woman; aged 19-60; **Fig. 1**, **Table 1**) were tested post-operatively at the Bethel Epilepsy Center (Bielefeld, Germany). The time between surgery and testing varied between one and six years (**Table 1**). In two patients (EM and BT), surgery did not result in a complete callosal transection; they are hereafter referred to as *partial splits*. In patient EM, the surgery was stopped after sectioning the anterior third of the CC; patient BT was spared approximately 1 cm of splenium. The remaining four patients (HY, DT, TJ, and LJ) were completely sectioned and are hereafter referred to as *full splits*. All patients have intact anterior and posterior commissures regardless of the extent of the callosal section.

**Figure 1.**
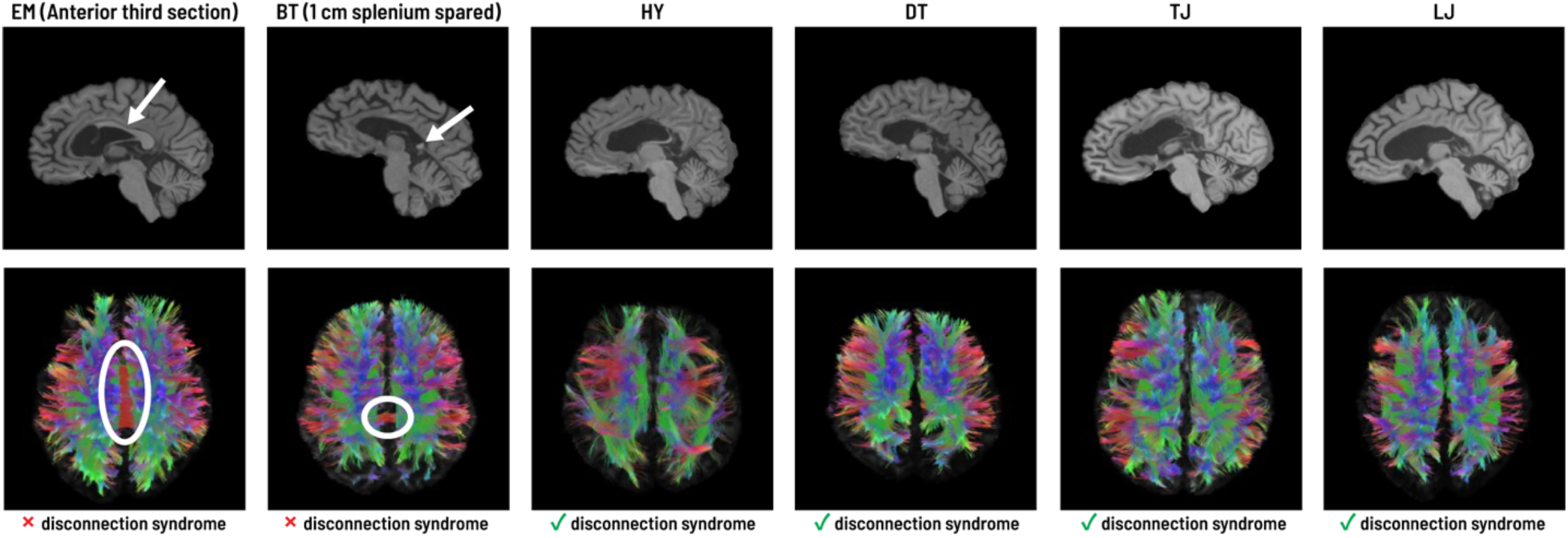
**Patient anatomical scans**. Post-operative *T*1 and diffusion MRI scans collected for all six patients at time of testing. White arrows/ellipses for patients EM and BT highlight intact CC (i.e., partial splits). The remaining four patients were fully-sectioned. Note that a series of clinical observations suggested behavioral disconnection syndromes in all full splits, but not in partial splits.

**Table 1.** General overview of callosotomy patients.

| Patient | Age at surgery | Age at testing | Sex | Handedness | Extent of callosotomy | Disconnection syndrome |
| --- | --- | --- | --- | --- | --- | --- |
| EM | 17 | 19 | M | Right | Partial | Absent |
| BT | 22 | 28 | M | Right | Partial | Absent |
| HY | 21 | 22 | M | Right | Complete | Present |
| DT | 15 | 20 | M | Right | Complete | Present |
| TJ | 49 | 52 | F | Right | Complete | Present |
| LJ | 57 | 60 | M | Right | Complete | Present |

Callosotomy is not commonly associated with post-operative neuropsychiatric or neurological disorders. Rather, split-brain patients often demonstrate *disconnection syndromes*, such that they appear to have two hemispheres operating independently, unable to communicate information from one half of the brain to the other^13,14,15^. Here, behavioral disconnection syndromes were clinically-observed using simple, lateralized bedside testing protocols. Comprehensive descriptions of these tests are described elsewhere^16^. In brief, all four complete callosotomy patients clearly indicated behavioral disconnections. Partial splits, in contrast, did not show *any* signs of disconnection syndromes—including, remarkably, patient BT (with a mere 1 cm of splenium). These findings contradict our classical understanding of callosal topography, which posits functionally-specific deficits depending upon which portions of the CC were severed. Consequently, a small proportion of posterior callosal fibers may be sufficient to support information integration across large-scale inter-hemispheric brain networks.

## Inter-hemispheric functional connectivity after callosotomy

All patients underwent resting-state functional magnetic resonance imaging without sedation, affording whole-brain analyses of resting-state functional connectivity (FC) and quantification of inter-hemispheric FC (IFC). To benchmark patients against ‘neurotypical’ patterns of IFC, we used data from the Human Connectome Project 100 Unrelated Subjects (HCP100) cohort. All HCP100 subjects and callosotomy patients underwent an identical preprocessing pipeline, including conservative corrections for mitigating head motion artifacts^20,21^ (see **Extended Data** Fig. 1 for a quantitative description of patient head motion). Whole-brain FC in the range of 0.01-0.08 Hz was assessed by parcellating the brain into 400 cortical regions and 32 subcortical regions (according to the Schaefer^17,18^ and Tian^19^ atlases, respectively; **Fig. 2a**) and correlating all pairs of regional time series (**Figs. 2b-c**).

**Figure 2.**
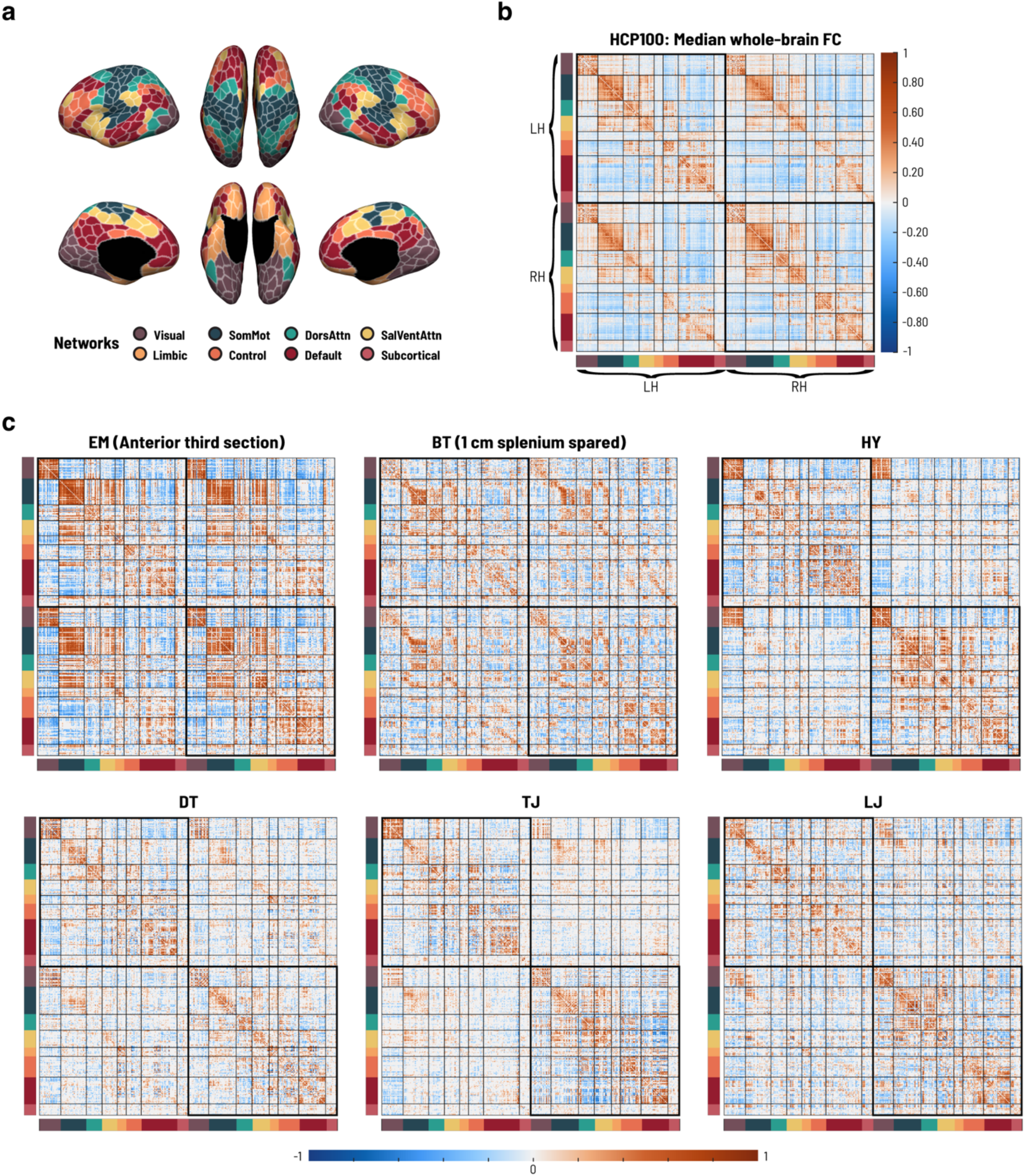
Inter-hemispheric functional connectivity is broadly-diminished in full (but not partial) splits. a, Whole-brain functional parcellation comprised of seven canonical resting-state networks^17,18^ and subcortical nodes^19^. **b,** A standard reference for neurotypical whole-brain functional connectivity (FC), given as the median over HCP100 subjects. FC was estimated by correlating low-frequency brain activity (0.01-0.08 Hz) between all pairs of regions in **(a)**. The large boxes outlining the upper left and lower right quadrants denote *intra-hemispheric* FC; off-diagonal quadrants thus capture *inter-hemispheric* FC (IFC). **c,** Whole-brain FC estimates for each callosotomy patient, thresholded at *p* < .001 (uncorrected for multiple comparisons). Typical patterns of IFC were largely abolished in full splits (HY, DT, TJ, LJ), while partial splits (EM, BT) were more similar to healthy controls. Abbreviations: SomMot, SomatoMotor Network; DorsAttn, Dorsal Attention Network; SalVentAttn, Salience/Ventral Attention Network; LH, Left Hemisphere; RH, Right Hemisphere.

All full splits demonstrated marked reductions in IFC relative to both neurotypical controls and partial splits (**Figs. 2b-c**). Spatial patterns of intra-hemispheric FC, by contrast, were generally better-matched to controls (**Extended Data** Fig. 2), suggesting that callosotomy primarily impaired inter-hemispheric functional coupling with full splits suffering the greatest disruptions. Curiously, however, while gross IFC was dissolved in full splits, inspection of individual FC matrices suggested that visual network connectivity was partially preserved (**Fig. 2c**). This may be due to synchronized perceptual inputs (scans were acquired with eyes open), driving *apparent* IFC despite the lack of transcallosal connectivity between visual regions and evidence for behavioral disconnections. We therefore sought to assess differences in the magnitude of IFC between functional homologues to determine whether any preserved IFC was unique to sensory systems. Here we computed a series of *network laterality indices*, where a value of zero indicates a perfect balance between inter- and intra-hemispheric FC, and values closer to -1 or 1 indicate relatively more inter-or intra-hemispheric FC, respectively. Note that we focus here on the distribution of positively-coupled edges (anti-correlated edges are shown in **Extended Data** Fig. 3).

Consistent with observed behavioral disconnections, all full splits demonstrated more lateralized patterns of FC relative to healthy controls and partial splits, who typically fell within the control distributions (**Fig. 3**). This was evident in both higher-order, transmodal systems (e.g., the default mode and dorsal attention networks) and in somatosensory networks (with the exception of patient TJ who fell within the upper tail of the control distribution in the somatomotor network). Patient EM was more lateralized across frontoparietal control nodes, which include a number of regions likely to be structurally-disconnected given the extent of their callosotomy; however, BT generally showed degrees of network bilaterality equivalent to healthy adults, despite having a mere fraction of splenium intact. Finally, full splits appeared to show conserved bilateral FC across limbic regions like the orbitofrontal cortex and temporal poles, which are known to be structurally coupled through the preserved anterior commissure as well as the CC^22,23,24^. In sum, these results highlight the dependence of inter-hemispheric networks on CC connectivity—yet, neurotypical levels of functional integration were possible even when only the splenium was left intact.

**Figure 3.**
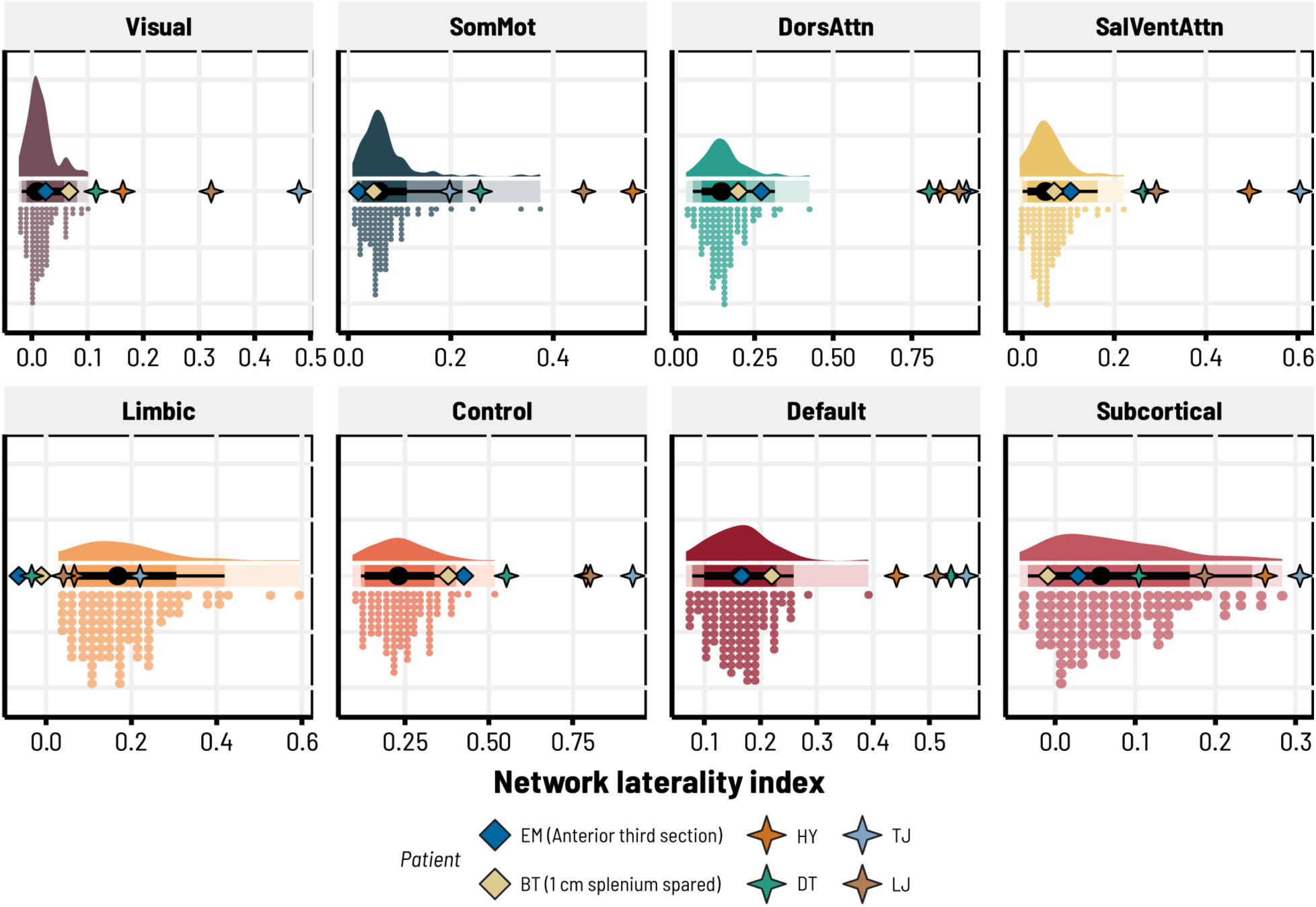
Full callosotomy is associated with more lateralized functional connectivity. Network laterality indices across seven canonical resting-state systems^17,18^ and subcortical regions^19^. Values closer to zero indicate an equal balance between intra-and inter-hemispheric FC, while values closer to one indicate more intra-hemispheric FC relative to inter-hemispheric FC. Note that each subplot captures a unique range of values, given differences in variability observed across neurotypical adults (colored distributions), patients, and networks. Full splits generally demonstrated more strongly-lateralized FC compared to partials, who fell within the control distributions. Point intervals for each control distribution give the HCP100 median (black dot) and the 80-95% quantiles around the median (thick black lines and thin black lines, respectively). Abbreviations: SomMot, SomatoMotor Network; DorsAttn, Dorsal Attention Network; SalVentAttn, Salience/Ventral Attention Network.

## Lateralized expressions of functional modules

To further elucidate whole-brain differences in functional network topology—agnostic to any *a priori* assignment of individual brain regions to canonical functional systems—we employed data-driven community detection techniques to cluster whole-brain patterns of FC into functional *modules* (**Fig. 4**). The modular organization of both structural and functional brain networks is thought to enable specialized information processing while retaining the ability to efficiently integrate information between disparate neural systems^25,26^. Thus, we hypothesized that the breakdown of IFC in full splits would produce lateralized network topologies, predominantly self-organized within one hemisphere or the other.

**Figure 4.**
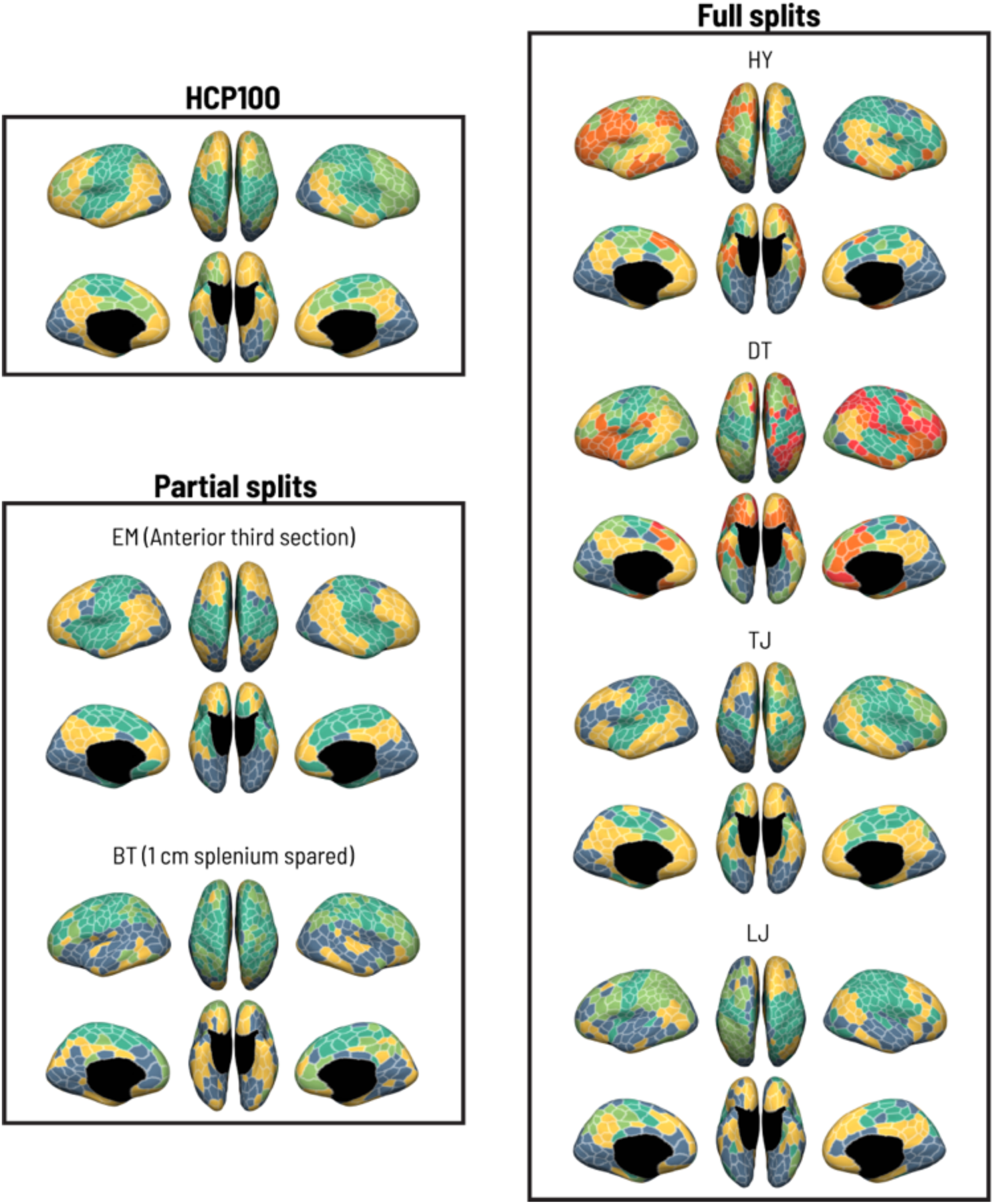
Modular decomposition of whole-brain functional connectivity. Spatial maps of optimal, consensus community assignments derived via data-driven modularity maximization at a default level of resolution (γ = 1). Regional coloring indicates assignment to a given functional module. Full splits generally exhibited more lateralized clustering of regions into modules relative to partial splits and healthy adult controls (see Fig. 5 for quantification of module laterality).

Here we computed laterality indices for the estimated partitions according to prior work^27^: where a value of zero, for example, indicates that all functional modules are perfectly bilateral (i.e., the constituent brain regions belonging to each community are equally-balanced between the left and right hemispheres), and values closer to one indicate progressively more lateralized topologies. Importantly, we note that modularity maximization is sensitive to spatial scale, subject to a *resolution parameter*, γ^28^. We describe results using a ‘default’ γ = 1 below but provide further consideration of these issues in the **Methods** and **Extended Data** Fig. 4.

As predicted, we observed much greater degrees of modular laterality in all full splits (**Fig. 5**). Communities with bihemispheric representations primarily spanned visual and somatomotor regions—reflecting a mechanism through which synchronized, external sensory inputs can drive apparent functional integration. With respect to partial splits, patient EM with the most CC showed bilateral patterns of network organization mirroring neurotypical controls. Patient BT also fell within the control distribution—however, they showed a strikingly different pattern of modularity (**Fig. 4**). Here, functional communities appeared to dissociate along the Sylvian fissure, with dorsal modules and ventral modules spanning the anteroposterior axis of the brain. This individual had the longest gap between surgery and testing (∼6 years), which may have allowed for widespread network reorganization affording optimal utilization of the minimal callosal fibers left intact. These findings underscore the importance of transcallosal connections for the emergence of coherent inter-hemispheric systems, while further highlighting a role for sensory inputs enabling apparent preservation of bilateral sensorimotor modules.

**Figure 5.**
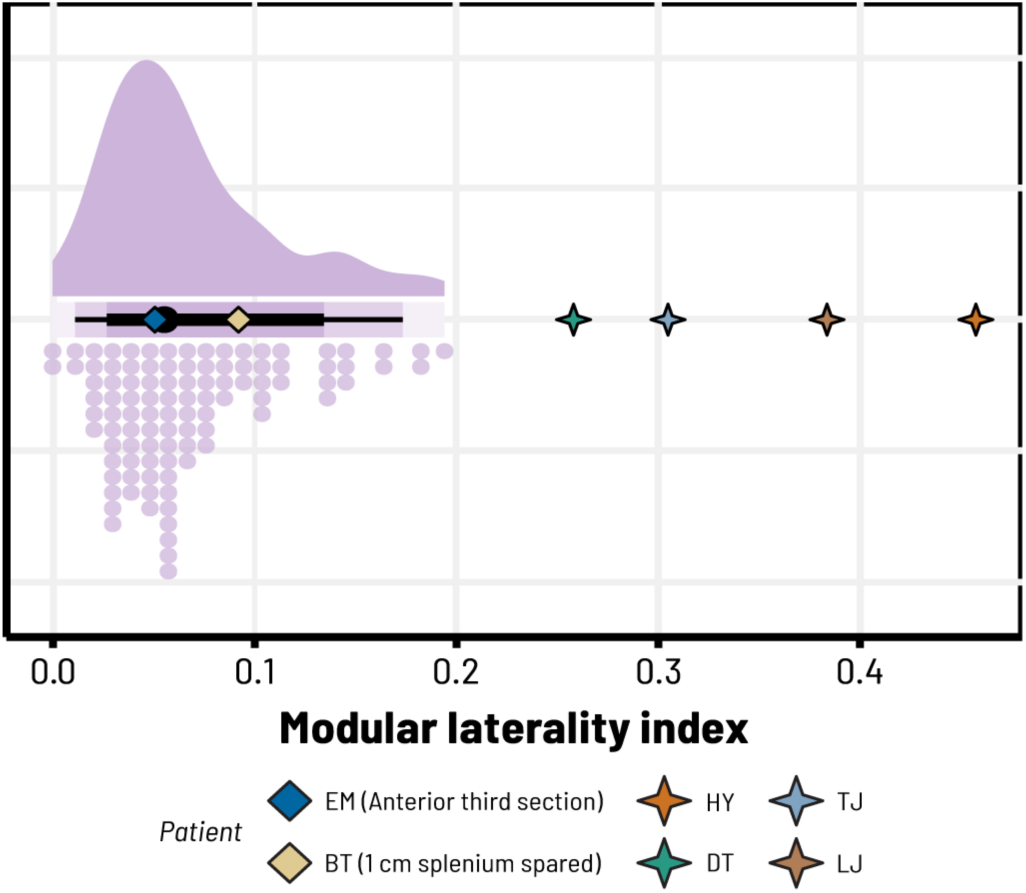
**Functional modules are more strongly-lateralized in full splits**. The modular laterality index quantifies the extent to which functional communities exhibit bihemispheric representations (values closer to zero) or are restricted to one hemisphere or the other (values closer to one). At a default level of resolution (γ = 1), partial splits demonstrated bilateral network topologies similar to healthy controls, whereas full splits were consistently more lateralized. Point intervals under the control distribution give the HCP100 median (black dot) and the 80-95% quantiles around the median (thick black lines and thin black lines, respectively).

## Inter-hemispheric synchrony in network dynamics

The modular organization of the brain plays a critical role in governing large-scale functional dynamics, such that structural architectures orchestrate widespread co-fluctuations in whole-brain activity over time^29^. Such patterns of coordinated network dynamics have recently been elucidated using *edge time series* (ETS): in brief, the ETS approach constitutes a temporal unwrapping of the zero-order Pearson correlation between any given pair of brain regions, capturing moment-to-moment changes in the extent to which regional pairs are co-fluctuating in activity^30^. Taking the root-sum-of-squares (RSS) across all ETS at each timepoint yields summary estimates of multivariate co-fluctuations in functional network dynamics over the course of a scan^31^ (**Fig. 6a**).

**Figure 6.**
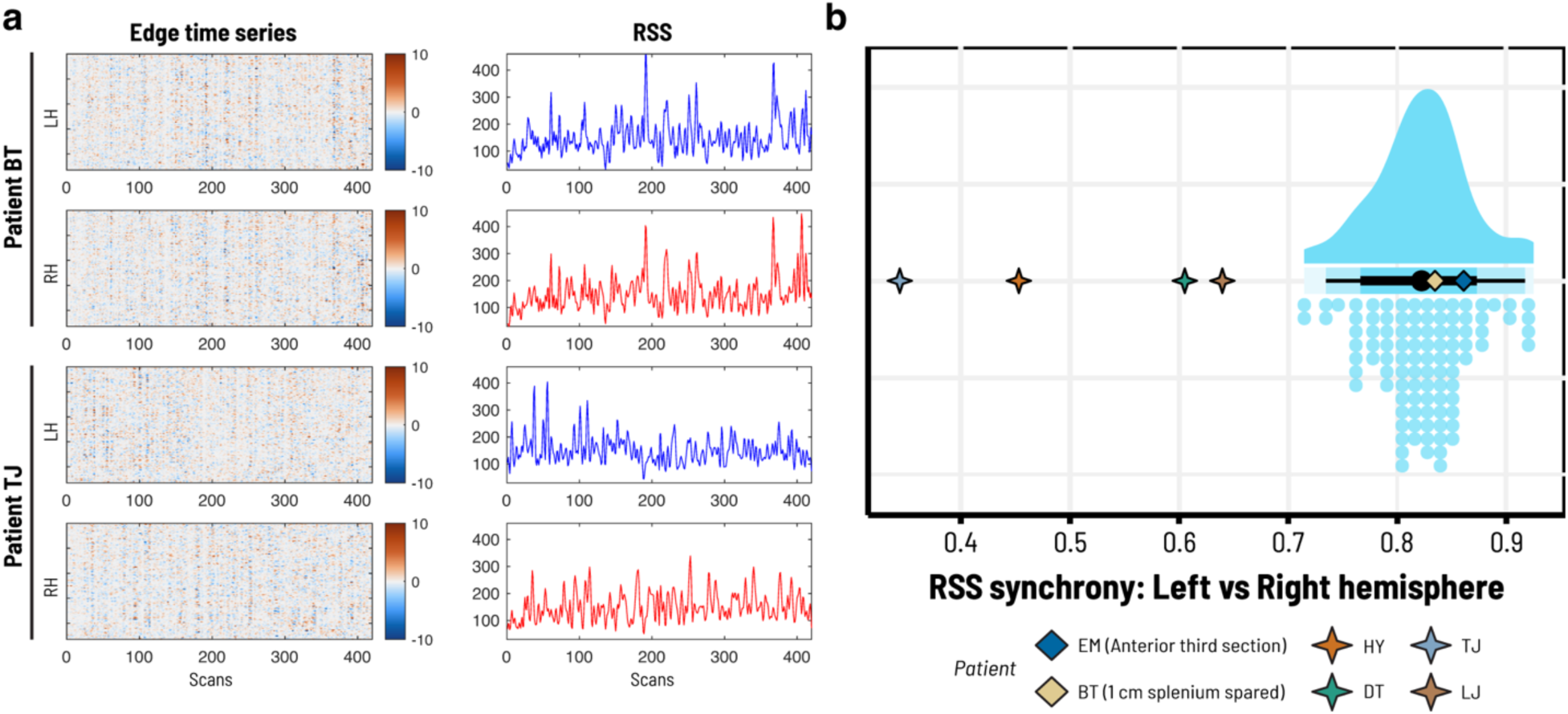
**Large-scale functional network dynamics are desynchronized in full splits**. **a,** An illustration of edge time series in the left and right hemispheres for patient BT (a partial split with 1 cm of intact splenium) and patient TJ (a full split). By taking the root-sum-of-squares (RSS) across all edges at each timepoint, we derived a summary estimate of the total amplitude of co-fluctuating brain activity. Note the bursty, high-amplitude co-fluctuations in each hemisphere that often coincide in time for BT, but not in TJ. **b,** All four full splits exhibited reduced temporal synchrony (i.e., temporal correlations) between left/right hemisphere RSS time courses. Partial splits, by contrast, fell within the range of healthy adult controls. Point intervals under the control distribution give the HCP100 median (black dot) and the 80-95% quantiles around the median (thick black lines and thin black lines, respectively).

Here we sought to test the hypothesis that structural connectivity via the CC is necessary for large-scale coordination of function. To that end, we separated edges within the left and right hemispheres and derived an RSS time course for each. We then computed correlations between left and right RSS to assess the degree to which dynamic co-fluctuations in brain function were synchronized across hemispheres. Among neurotypical controls, we observed tight temporal coupling between left and right hemisphere dynamics (Median Spearman *r* = 0.82, *MAD* = 0.08). In contrast, all four full splits demonstrated significant reductions in inter-hemispheric synchrony, suggesting that network dynamics were more functionally-independent (**Fig. 6b**; see **Fig. S4** for individual networks). This is consistent with the overall attenuation of inter-hemispheric FC and further indicates that brief moments of high-amplitude co-fluctuations (which normally encompass regions in both hemispheres and are thought to be structurally-mediated^29^) occur separately in each. Notably, both partial splits fell within the standard range of the HCP subjects. These results therefore strongly suggest a mechanistic role for CC structural connections in supporting widespread integration of inter-hemispheric network dynamics. This may be achieved with a mere fraction of intact fibers through the splenium and thus might be largely-independent of the presence of direct cortico-cortical connections.

## Discussion

Distinct disruptions of inter-hemispheric functional connectivity were observed following complete versus partial transection of the corpus callosum. Full splits demonstrated more lateralized network topologies and desynchronized network dynamics, aligning with classical behavioral disconnection syndromes. Given that all patients had intact anterior and posterior commissures (along with other subcortical routes of inter-hemispheric communication), these findings underscore the notion that transcallosal connections are the essential pathways through which large-scale inter-hemispheric functional networks emerge. Contrary to our expectations, however, inter-hemispheric functional coupling among partial splits remained intact. These patients also showed *no* signs of behavioral disconnection, even when the vast majority of the callosum was severed^16^. Thus, a fraction of callosal fibers in the splenium may be sufficient to orchestrate inter-hemispheric functional synchrony and integrate behavior. This poses a remarkable challenge to our classical topographic models of callosal connectivity and function, motivating a novel view on inter-hemispheric structure-function relationships: namely, that robust inter-hemispheric synchrony and information transfer can be achieved via polysynaptic connectivity through a relatively small proportion of posterior transcallosal connections, rather than direct structural connectivity between functional homologues.

## Complete callosotomy and structure-function relationships

Split-brain patients present a unique opportunity to draw causal inferences regarding the role of transcallosal structural connections and the extent to which inter-hemispheric functional synchrony depends upon such connections. Previous investigations of pediatric callosotomy patients have provided an initial perspective on this question^9,10,11,12^. However, these patients often suffer from severe cognitive and developmental deficits, requiring scans to be completed under sedation—which reduces dynamical complexity in brain activity and produces functional connectomes more rigidly tied to direct anatomical pathways^32^. It is further known that functional and structural brain networks undergo profound developmental changes early in life^33,34,35,36^. In other words, neural plasticity in these young individuals may still afford the long-term establishment of alternate inter-hemispheric pathways (perhaps akin to callosal agenesis patients^37^), and indeed, it is well-documented that disconnection syndromes in pediatric patients resolve over time^38,39^. This likely accounts for differential reports of recovery in inter-hemispheric FC among these individuals^9,11,12^.

The adult human brain also retains neuroplastic potential^40^. However, given the enormity of the structural disconnection, adult callosotomy patients may lack the capacity to adapt their functional architectures in such a way that allows complete recovery of communication between disconnected brain regions. Our results clearly underscore this hypothesis. Whole-brain patterns of FC appeared highly lateralized, both with respect to canonical network definitions (**Figs. 2-3**) and the topological organization of functional modules (**Figs. 4-5**). Furthermore, we observed considerable reductions in the extent to which multivariate co-fluctuations in activity were synchronized between hemispheres (**Fig. 6**), underscoring a role for the CC in orchestrating whole-brain network dynamics. Importantly, however, there were some exceptions to these effects, which follow anatomical predictions. Limbic and subcortical regions—the former of which are known to be connected through the anterior commissure (AC) as well as the CC^22,23,24^—showed conserved bilateral FC (**Fig. 3**). Nevertheless, non-callosal pathways were insufficient to support widespread inter-hemispheric synchrony, contrary to previous proposals both in humans^41,42,43^ and nonhuman primates (where AC fibers are sufficient to sustain inter-hemispheric FC)^44^. Finally, visual and sensorimotor areas also appeared to remain somewhat integrated. We cannot fully discount the possibility that subcortical information transfer contributed to this preserved synchrony; however, an alternative hypothesis may lie in the fact that visual and somatosensory inputs to each hemisphere were synchronized during the resting-state scan—producing *apparent* functional coherence despite observed behavioral disconnections.

## Preserved inter-hemispheric integration in partial splits

While our observations in full splits clearly emphasize the importance of CC connectivity in orchestrating inter-hemispheric synchrony, partial splits revealed something much more surprising. We predicted that partial splits would show specific deficits in inter-hemispheric FC along an anterior-to-posterior gradient localized to the extent of the callosal transection, following both previous characterizations of CC topography^7,8^ and prior behavioral studies^45,46,47,48^. However, both partial splits consistently demonstrated levels of inter-hemispheric integration in the range of neurotypical controls (**Figs. 2-6**). This was even observed for patient BT, in whom only ∼1 cm of posterior callosal fibers were spared. It has become increasingly-evident that structure-function relationships are highly complex and generally do not reflect simple, one-to-one interactions between directly-coupled areas^49,50,51^. Still, these remarkable results imply that the functional topography of the CC may be much more flexible than previously-imagined. In particular, there may be a critical threshold at which full functional integration can be sustained given a sufficient proportion of intact fibers—even if the vast majority of the callosum has been severed.

An intriguing extension of this hypothesis is that the *location* of the spared callosal fibers may be critical. Previous cytological work has shown that anterior portions of the CC are primarily populated by thin fibers, while fibers in the tail locally-vary in diameter^52,53,54^. Such differences in fiber size and myelination along the CC have been associated with differences in conduction velocity^54^ and, when considered alongside the topographic organization of the CC, may account for the ‘fast’ versus ‘slow’ action of primary sensorimotor and association cortices, respectively. Spared posterior fibers as in BT could therefore be more beneficial than spared anterior fibers. However, from a sheer bandwidth perspective, it is difficult to imagine how such a small proportion of fibers could account for widespread integration of function and cognitive processing.

It appears likely that functional systems in BT dramatically reorganized in the six years following surgery. In brains that develop normally with the support of the CC, multisensory integration zones in parietal cortex may serve as inter-hemispheric hubs^55,56^. By taking advantage of spared posterior callosal pathways connecting these regions, the processing of various intra-hemispheric brain regions can still be integrated and exchanged along polysynaptic routes running between hemispheres. The striking differences in functional network topology observed in BT, with large modules spanning the anterior-to-posterior axis of the brain (**Fig. 4**), are one potential consequence of this reorganization, reflecting greater functional dependencies between frontal and posterior areas affording more optimal usage of the spared splenial fibers. The fundamental importance of these posterior inter-hemispheric connections also aligns with and may further inform neuroscientific theories of consciousness^57,58^. While our results are not suited to evaluate or endorse any particular theory, it is nevertheless curious that the temporal-parietal-occipital ‘hot zone’ of regions recently proposed to maximize the singular ‘here-and-now’ *qualia* of consciousness^57,59^ are likely dependent (in part) upon CC connections through the splenium. This may explain why BT maintains an integrated conscious experience while full splits demonstrate inter-hemispheric disintegration^16^.

## Limitations

We acknowledge the small number of patients included in this study, each of whom have limited post-operative data. While previous pediatric studies have recruited larger samples, the availability of patients who underwent callosotomy in adulthood is extremely scarce. It is often difficult to draw robust and reliable statistical inferences from neuroimaging data acquired on such small samples. However, the marked differences observed between full splits and healthy controls suggest that we are not parsing trivial effects on the scale of traditional neuroimaging—rather, they reflect outcomes drawn from fundamentally unique sampling distributions. Furthermore, relative to other lesion studies (often hampered by variability in spatial localization/extent), callosotomy is a highly-selective intervention. By severing a single white matter structure in the brain without any damage to gray matter, callosotomy may afford more homogeneous outcomes. It will be critical for future studies to include larger samples with more imaging data to better characterize the extent of neural variability in callosotomy patients, especially as network architectures reorganize over time.

## Conclusion

Following more than a half-century of behavioral neuroscience, split-brain patients continue to provide startling insights into the fundamental nature of the human brain. The rise of network neuroscience has further transformed our understanding of large-scale brain systems, tracing the pathways relating brain structure to function and behavior. Our findings reveal a unique type of criticality in brain function—such that a vanishingly-small proportion of posterior callosal fibers may be sufficient to support the integration of inter-hemispheric functional brain networks and unified cognitive processing. It has long been celebrated that the human brain is a remarkably adaptable complex system. These findings challenge us to rethink our conventional assumptions about structure-function relationships and the means through which the brain enables the mind.

## Online Methods

### Patient recruitment

Six adult callosotomy patients (5 men, 1 woman; aged 19-60; **Fig. 1**, **Table 1**) were recruited during routine post-operative follow-up visits to the Bethel Epilepsy Center (Bielefeld, Germany). The study was approved by the Ethics Committee of the University of Münster, Germany (2021–523-f-S). The patients or their legal representatives provided written informed consent.

## Callosotomy procedure

Callosotomies were performed with the assistance of intraoperative neuronavigation, with patients in a supine position and the head inclined by roughly 30° (fixed in the Mayfield clamp). A straight skin incision was made running slightly next to the midline on the non-language dominant hemisphere. Craniotomy was then performed by making one drill hole in the midline at approximately the level of the precentral sulcus. A bone flap was sawn out with an outline of 3 cm x 5 cm. The dura was incised longitudinally and then opened with two wings under microneurosurgical conditions—access to the inter-hemispheric fissure was successively created while preserving the cortical bridging veins. Next, using a self-retaining retractor to carefully shift the frontal lobe, the level of the corpus callosum was gradually exposed under sharp dissection with micro scissors.

Dissection is often made considerably more difficult by the course of the pericallosal arteries, which run parallel to the corpus callosum but are often tightly intertwined and twisted in the inter-hemispheric fissure. In cases where the risk of vascular damage is too great, the surgery may be halted short of a full disconnection (resulting in partial splits). However, under most conditions, the inter-hemispheric fissure may be opened along the entire length of the knee of the corpus callosum. The corpus callosum was then perforated into the lateral ventricle of the ipsilateral hemisphere. Next, following the pericallosal artery and the anterior cerebral artery down to the frontal base, the knee of the callosum was split in the midline, continuing backwards to about the level of the foramen of Monroi. From this point, it is critical to protect the fornix of the language-dominant hemisphere: to mitigate the risk of postoperative functional impairment, we carefully ensure that the self-retaining retractor is not placed on the precentral gyrus within the inter-hemispheric fissure. The disconnection can now be carried out completely into the splenium—the severing of the splenium was performed subpially to protect all the cisternal vessels. Extensive hemostasis is not usually necessary, and thus a ventricular drainage is not used. To conclude, the dura mater was closed and the bone flap reinserted and fixed with titanium plates and screws. Finally, a subgaleal drainage was inserted and the skin closed in two layers.

## Healthy adult controls

We used open-access data from the Human Connectome Project 100 Unrelated Subjects (HCP100) cohort (46 men, 54 women; *M*_Age_ = 29.11, *SD* = 3.68) as a robust benchmark for various network features estimated in callosotomy patients. Participant recruitment and imaging procedures have previously been detailed elsewhere^60,61^. In brief, data were collected over two sessions on a custom 3T Siemens Skyra with a 32-channel head coil. *T*_1_-weighted structural scans were acquired using a 3D magnetization prepared rapid gradient echo (MPRAGE) sequence (*TR* = 2400 ms, *TE* = 2.14 ms, *TI* = 1000 ms, flip angle = 8°, voxel size = 0.7 mm^3^). Resting-state fMRI was collected with eyes open over four ∼14-minute scans, using a multiband echo-planar imaging (EPI) sequence sensitive to the blood-oxygen-level-dependent (BOLD) contrast (*TR* = 720 ms, *TE* = 33.1 ms, flip angle = 52°, voxel size = 2 mm^3^, multiband factor = 8). All participants gave written informed consent for a protocol approved by the Washington University Institutional Review Board. All imaging data were downloaded in their raw (unprocessed) state.

## Patient neuroimaging procedures

Patient data were collected on a 3T Siemens Vida at the Bethel Epilepsy Center using a 32-channel head coil. *T*_1_-weighted structural scans were acquired using a 3D MPRAGE sequence (*TR* = 1900 ms, *TE* = 2.47 ms, *TI* = 900 ms, flip angle = 8°, voxel size = 0.8 mm^3^). Resting-state fMRI was collected with eyes open using a single-shot EPI sequence sensitive to the BOLD contrast (*TR* = 2000 ms, *TE* = 30 ms, flip angle = 75°, voxel size = 3 x 3 x 3.30 mm, iPAT/GRAPPA factor = 2).

We note that each patient varied somewhat in the amount of resting-state data obtained (from ∼8-20 minutes total) due to differences in clinical scheduling and other imaging protocols not relevant to the current analysis. This information is summarized in **Extended Data** Fig. 1.

## fMRI preprocessing

All data (from both patients and healthy controls) were run through a unified preprocessing pipeline. Initial preprocessing relied on functions from the Statistical Parametric Mapping 12 software (SPM12, Wellcome Trust Centre for Neuroimaging, London) in Matlab and Advanced Normalization Tools (ANTs)^62^. Resting-state fMRI data were realigned and unwarped to correct for head motion and motion x susceptibility artifact as well as slice-time corrected. Anatomical images were denoised (ANTs’ DenoiseImage with a spatially-adaptive Gaussian filter) and submitted to ANTs’ cortical thickness pipeline, which includes N4 bias field correction and high-quality image segmentation and brain extraction^63^. For some patients, the brain extraction mask included extracerebral components: this was corrected by manually eroding the extraction mask and re-applying it to the anatomical scan when necessary. Image coregistration parameters were subsequently estimated between an N3 bias-corrected mean BOLD reference and the skull-stripped *T*_1_ image via antsRegistration (including both linear and nonlinear transformations to minimize residual signal deformations, cf. ‘SyN distortion correction’ as implemented by the widely-used fMRIPrep and QSIPrep pipelines).

Subsequent spatial normalization to a standard stereotaxic space presented a nontrivial problem for patient data—in the absence of a corpus callosum, generic registration routines (with both linear and nonlinear stages) will frequently pull structures such as the cingulate gyrus into the space occupied by the CC in the MNI template image, which misaligns brain regions relative to their location in our functional parcellation. We mitigated this problem in two ways. First, we implemented cost-function masking during the nonlinear step to minimize the influence of callosal regions that were transected by the surgery: these masks were specified in the fixed image space (i.e., the MNI152NLin6Asym template, as provided by the FMRIB Software Library, FSL^64,65^) because we lacked pre-operative data for each patient, and CC segments were defined by the JHU-DTI white matter atlas^66,67^ (also provided in FSL). For patient EM, the genu was masked out; for patient BT, all but the splenium was masked out; and for all full splits, the entire cross-section of the callosum was masked out. Second, in addition to these masks, we specified a series of registration metric objectives at the nonlinear step. As is typical for antsRegistration, we estimated similarity between the MNI template and *T*_1_ images using both ANTs’ neighborhood cross-correlation and Mattes mutual information similarity metrics. We also provided smoothed segmentation posteriors derived from the ANTs cortical thickness pipeline (six in total: white matter, cortical gray matter, CSF, deep/subcortical gray matter, brain stem, and cerebellum) and estimated similarity between the MNI template segments and patient segments using a mean-squares objective, with cortical white matter and CSF segments downweighted in importance relative to the other four segments (by a factor of one third). Thus, the nonlinear step was comprised of eight similarity metrics. The transformation was estimated using BSplineSyN to allow for smoother deformations without sacrificing accuracy.

Ultimately, we concatenated and applied the estimated coregistration and spatial normalization parameters to all fMRI data in a single interpolation via antsApplyTransforms, resampled into 2 mm^3^ isotropic voxels. Registration quality was visually assessed to ensure appropriate mapping to MNI space. Functional images were then smoothed using a 3D Gaussian kernel with FWHM = 5 mm^3^. Prior to functional connectivity estimation, several additional preprocessing steps were performed using in-house Matlab code: this included global signal scaling (median = 1000), linear detrending, and nuisance regression. The nuisance regression model followed a ‘36-parameter’ (36P) approach including the six motion regressors from realignment, the first eigenvariate of white matter signal, the first eigenvariate of CSF signal, and mean global signal—along with their squares, their first derivatives (via backwards-differentiation), and the squares of the derivatives. All nuisance regressors were detrended to match the BOLD data. We further augmented the design matrix of the nuisance model with a series of binary spike regressors: one for each image volume where the relative framewise displacement in head motion (accounting for both translation and rotation) was ≥ 0.50 mm. This ‘36P + spike regression’ model has importantly been shown to outperform alternative confound regression strategies (e.g., motion scrubbing and ICA-based methods) and provide the best control for motion artifact in functional connectivity analyses^20,21^. Patients generally showed no more head motion than HCP subjects; for reference, we provide a summary of head motion parameters and the frequency of motion spikes for each patient in **Extended Data** Fig. 1.

## Functional connectivity estimation and network laterality

Functional network nodes were defined by parcellating the brain into 400 cortical regions and 32 subcortical regions, according to the Schaefer^17^ and Tian^19^ atlases, respectively. We extracted a summary time course from each parcel by taking the first eigenvariate over time^68^. These regional time series were bandpass-filtered to the range of 0.01-0.08 Hz. We then constructed 432 x 432 FC matrices by estimating the zero-order Pearson correlation between all pairs of regional time series. For both HCP100 subjects and callosotomy patients with more than one resting-state scan, runs were concatenated prior to FC estimation.

We sought to quantify differences in inter-hemispheric functional connectivity (IFC) between callosotomy patients and healthy controls from a number of network neuroscience perspectives. Pursuant to this, we began with an initial, breadth-first assessment of IFC within each of the seven canonical resting-state networks defined by our cortical parcellation^17,18^, as well as regions spanning the subcortex^19^. Similarly to studies of functional lateralization in task-based fMRI, we derived a *network laterality index* as a simple means of summarizing and quantifying the *relative magnitude* of IFC as compared to the magnitude of intra-hemispheric FC. Each FC matrix was first thresholded at an uncorrected level of *p* < .001 to avoid overly-conservative and highly-nonlinear effects of multiple comparisons correction on the distribution of edge weights, while simultaneously minimizing the influence of weak (potentially spurious) connections. We then Fisher-*Z* transformed the thresholded FC matrices and, for each individual and for each network, estimated a laterality index, *LI*, as:

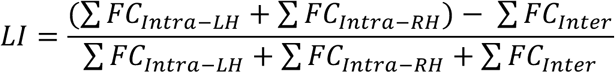

Intra-hemispheric connections within the left and right hemispheres are necessarily summed in the numerator given that there are, by definition, twice as many inter-hemispheric connections as each hemisphere alone. The result is a ratio on the interval of [-1, 1] that captures the extent to which each canonical resting-state network is predominantly connected intra-or inter-hemispherically: in the limit, a value of -1 would suggest that there is *no* intra-hemispheric FC and the network is solely comprised of inter-hemispheric connections, while a value of 1 would indicate the reverse. A value of 0 therefore represents a network that is perfectly balanced in the magnitude of intra-vs. inter-hemispheric connections. Note that the results presented in **Fig. 3** are with respect to *positively-correlated edges* only; we provide a comparison of anti-correlated edge lateralization in **Extended Data** Fig. 3.

## Community detection and modularity

While the above approach is useful for considering differences among canonical functional networks, these systems were defined in healthy adult brains—it is therefore perhaps dubious to assume that these networks follow a similar spatial organization in callosotomy patients. Data-driven community detection techniques afford assessments of functional network topology unbeholden to prior assumptions about specific brain regions and their canonical network memberships^26,69^. This is achieved via decomposition of FC matrices using *modularity maximization* algorithms, which group regions into *functional modules*: these represent regions that are generally more functionally-coupled to each other than they are connected to other regions. The optimization objective is subject to a *quality* metric, *Q*—unlike our previous laterality analysis (which required separating out positive and negative edges), this can be estimated over FC matrices comprised of both positive and negative correlations^70^. We performed this decomposition using the community_louvain function from the Brain Connectivity Toolbox (https://sites.google.com/site/bctnet/) with asymmetric treatment of negative edge weights^70,71^. Importantly, modularity maximization is sensitive to spatial scale, subject to a *resolution parameter*, γ^28^. Small values of γ typically yield coarse partitions with a small number of large communities; larger values of γ, by contrast, afford more fine-grained partitions. Because the solution of this decomposition is dependent upon one’s choice for the resolution parameter, we tested a range of values from γ = [0.10, 2] (in steps of 0.10). To further account for variability in community assignments (e.g., due to algorithmic stochasticity arising from random initializations), we estimated modularity 1000 times for each value of γ—optimal consensus partitions were obtained using the consensus_similarity function from the Network Community Toolbox (https://commdetect.weebly.com/).

In similar fashion to the network laterality indices described above, we derived a *module laterality index* for the resulting partitions according to prior work^27^. This metric is limited to the [0, 1] interval, where a value of 0, for example, indicates that all functional communities are perfectly bilateral (i.e., all of the constituent brain regions belonging to each community are equally-represented between the left and right hemispheres), and values closer to 1 indicate progressively more lateralized topologies. We note that in this case, more fine-grained partitions may be beneficial for assessing modular laterality—nevertheless, the results shown in **Figs. 4-5** reflect a ‘default’ γ = 1 and we further illustrate in **Extended Data** Fig. 4 that full splits rapidly diverge from controls even at smaller values of γ (while partial splits and healthy controls remain relatively consistent, regardless of γ).

## Inter-hemispheric network dynamics

Finally, we used edge time series (ETS)^30^ to assess the extent to which *many* brain regions in the left and right hemispheres dynamically co-fluctuate in synchrony. The ETS approach has been used to obtain moment-to-moment, time-resolved estimates of FC and characterize the nature of high-amplitude events^31^ that occur when multivariate BOLD signals simultaneously fluctuate together over brief intervals of time—these bursts of synchronized network dynamics are sufficient to recapitulate ‘static’ FC estimates over the course of a scan^31^, and importantly, they are independent of head motion^30^. Current evidence instead suggests these high-amplitude co-fluctuations are governed, in part, by the modular organization of the brain and its underlying structural architecture^29^. An ETS can be computed by taking a pair of BOLD time series from two regions, *Z*-scoring them, and taking the elementwise product of the signals at each timepoint—the mean of an ETS is equivalent to the zero-order Pearson correlation between the two time series, and thus ETS constitute a temporal-unwrapping of FC over the scan^30^.

Here, we first separated out all regions in the left hemisphere and all regions in the right hemisphere. We constructed a *T* x 23220 ETS matrix for each hemisphere, where *T* gives the number of timepoints and 23220 gives the total number of edges within each hemisphere. A summary time course quantifying the total amplitude of intra-hemispheric co-fluctuations at any given moment (i.e., across *all* edges) was obtained by taking the root-sum-of-squares (RSS) at each timepoint^31^. We then computed the Spearman correlation between the left hemisphere RSS and right hemisphere RSS time series to capture the extent to which network dynamics in each hemisphere were synchronized.

## Data and code availability

Patient data will be made available upon reasonable request to the corresponding authors; all relevant processing and analysis code will be uploaded to a Github repository.

## Acknowledgements

T.S., S.B., J.M.S., H.E.S., B.G., and M.B.M. were sponsored by the Army Research Office under contract W911NF-19-D-0001 for the Institute for Collaborative Biotechnologies. T.P. and L.J.V were funded by the German Research Foundation (CRC-1451, Project 431549029).

## Author contributions

T.S., S.B., T.P., H.E.S., J.L.H., A.R., B.G., C.G.B., O.S., M.S.G., L.J.V., and M.B.M. conceived and planned the experiments. T.P., V.M.W., J.L.H., A.R., F.G.W., C.G.B., and L.J.V. assisted with patient recruitment and were responsible for data collection. T.K. performed the callosotomy surgeries. T.S., J.M.S., O.S., and L.J.V. conceived the data analysis strategy; T.S., J.M.S., and O.S. contributed to analysis code and executed the data processing/analysis pipeline. T.S. wrote the manuscript with contributions from T.K., O.S., M.S.G., L.J.V., and M.B.M. The project was supervised by A.R., C.G.B., L.J.V., and M.B.M. Funding was secured by B.G., L.J.V., and M.B.M. All authors assisted with manuscript revision and approved the final submitted version. L.J.V. and M.B.M. contributed equally.

## Conflicts of Interest

The authors declare no competing interests.

---

**Extended Data Fig. 1.**
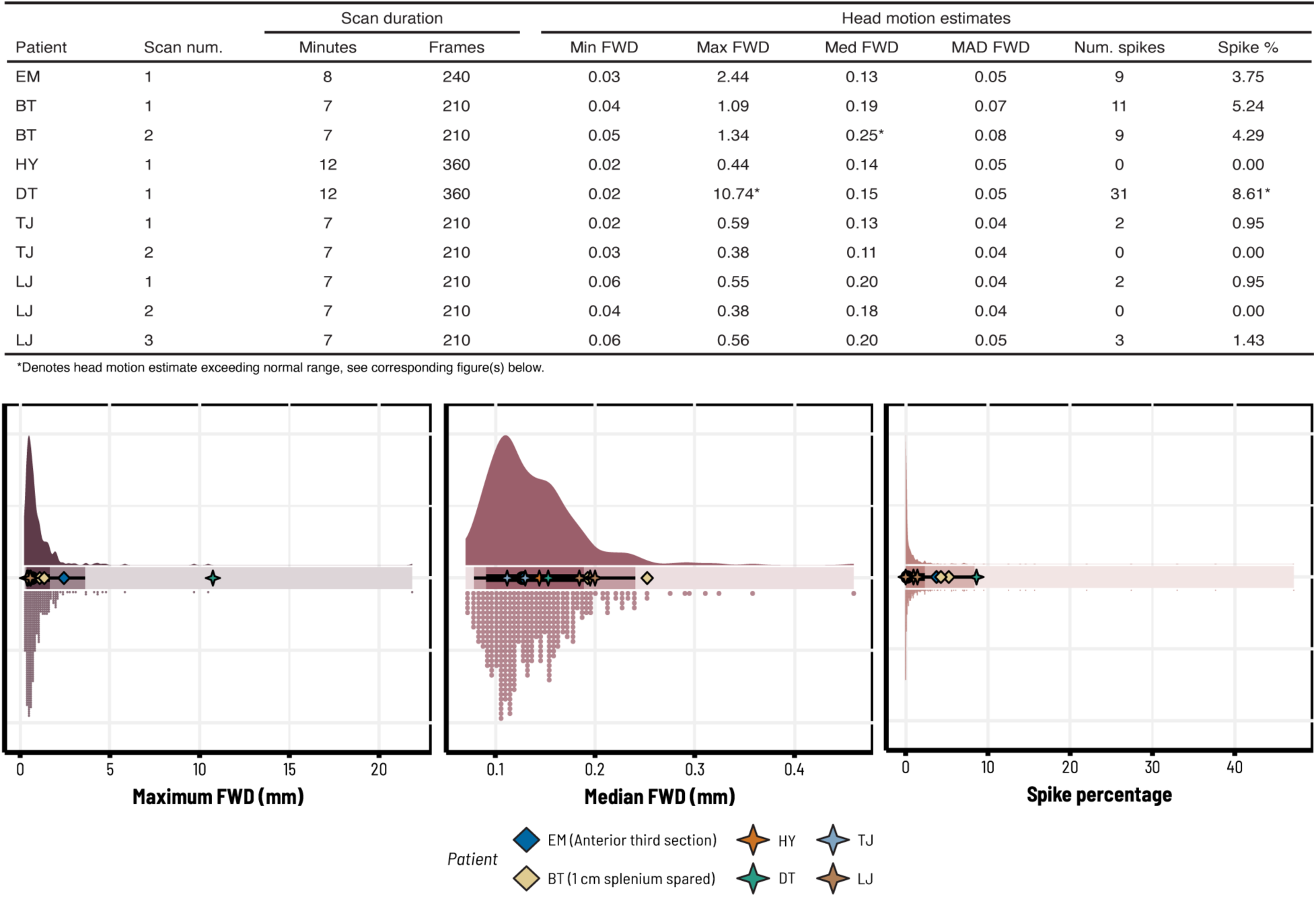
Scan times and head motion estimates across callosotomy patients. Due to differences in clinical scheduling and other imaging protocols not relevant to the current analysis, patients varied in resting-state fMRI scan time. Here we summarize these differences and provide various estimates of head motion for each scan. Motion parameters are given with respect to *framewise displacement* (FWD)—the amount of total head motion (in millimeters, considering both translational displacements and rotations) observed from one image volume to the next. Because FWD is a time series, we provide the minimum, maximum, and median FWD over each scan (with the median absolute deviation, *MAD*, over time). We also denote the number of motion spikes, defined as any frame with FWD ≥ 0.50 mm, and the percentage of frames contaminated with motion spikes—these were used to define individual spike regressors for nuisance signal regression^20,21^. Patients generally did not show any more head motion than healthy adult controls (see visualized distributions). Note that in rare cases where patients exceeded the range of controls in one metric, they were typically within the standard range of other metrics. Point intervals under the control distributions give the median (black dot) and the 80-95% quantiles around the median (thick black lines and thin black lines, respectively).

**Extended Data Fig. 2.**
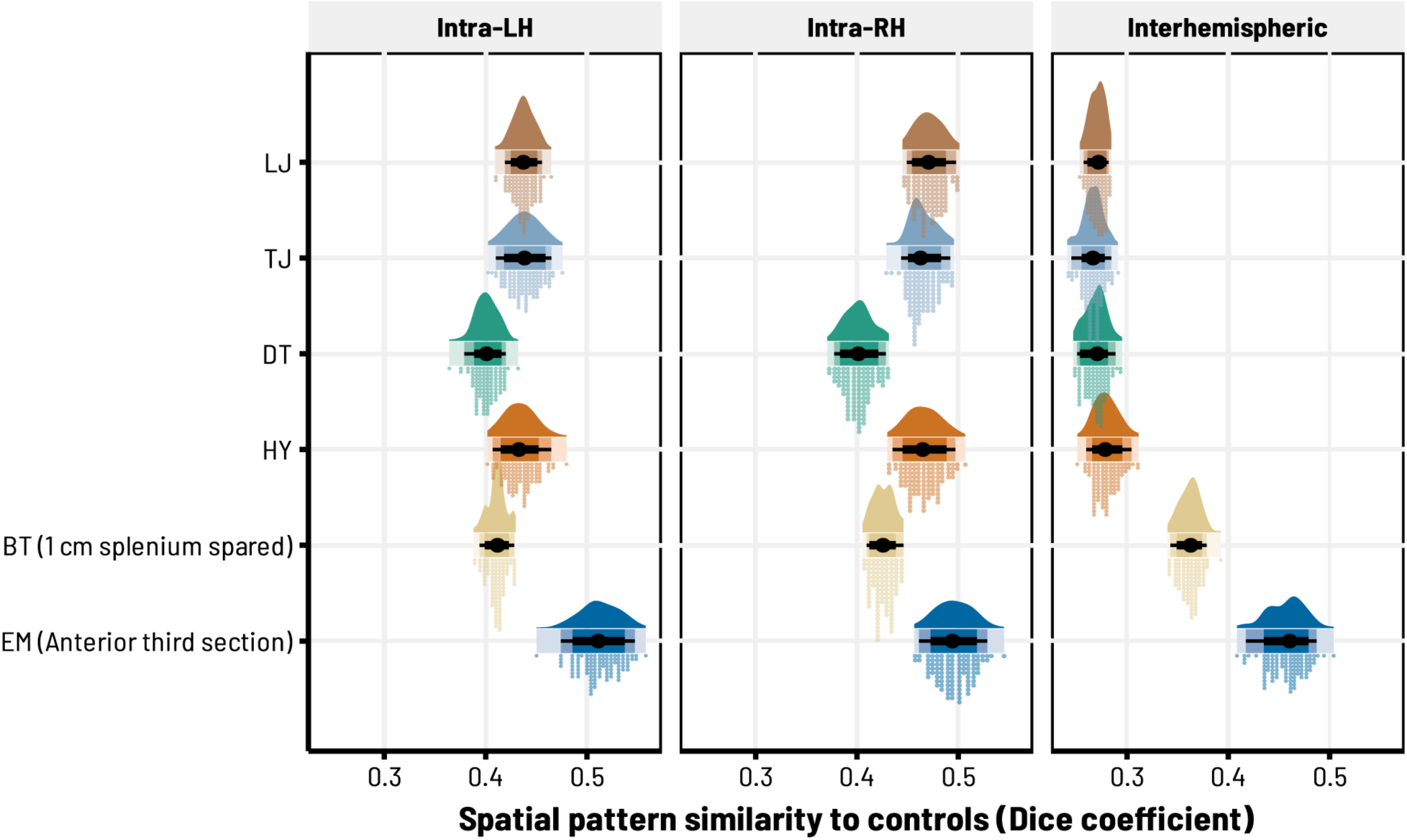
Multivariate similarity in spatial patterns of functional connectivity. As an initial estimate of the extent to which full vs. partial callosotomy disrupted intra-vs. inter-hemispheric functional connectivity (FC), we binarized FC matrices at a threshold of *z*(*r*) > 0 (where *z*(*r*) refers to the Fisher*-Z* transformed correlation coefficients for each network edge) and computed the Dice coefficient between each patient and each individual healthy control—this statistic ranges from [0,1] and quantifies the degree of similarity in two patterns of FC (liberally, whether the same edges show *any* FC > 0, regardless of the exact magnitude of those connections). The distributions for each patient thus show the range of similarity between a given patient and each control. Patient EM (with the most corpus callosum intact) is generally most similar to controls; full splits, on the other hand, show markedly dissimilar patterns of inter-hemispheric FC relative to intra-hemispheric FC. Point intervals under each distribution give the median Dice similarity (black dot) and the 80-95% quantiles around the median (thick black lines and thin black lines, respectively). Abbreviations: LH, Left Hemisphere; RH, Right Hemisphere.

**Extended Data Fig. 3.**
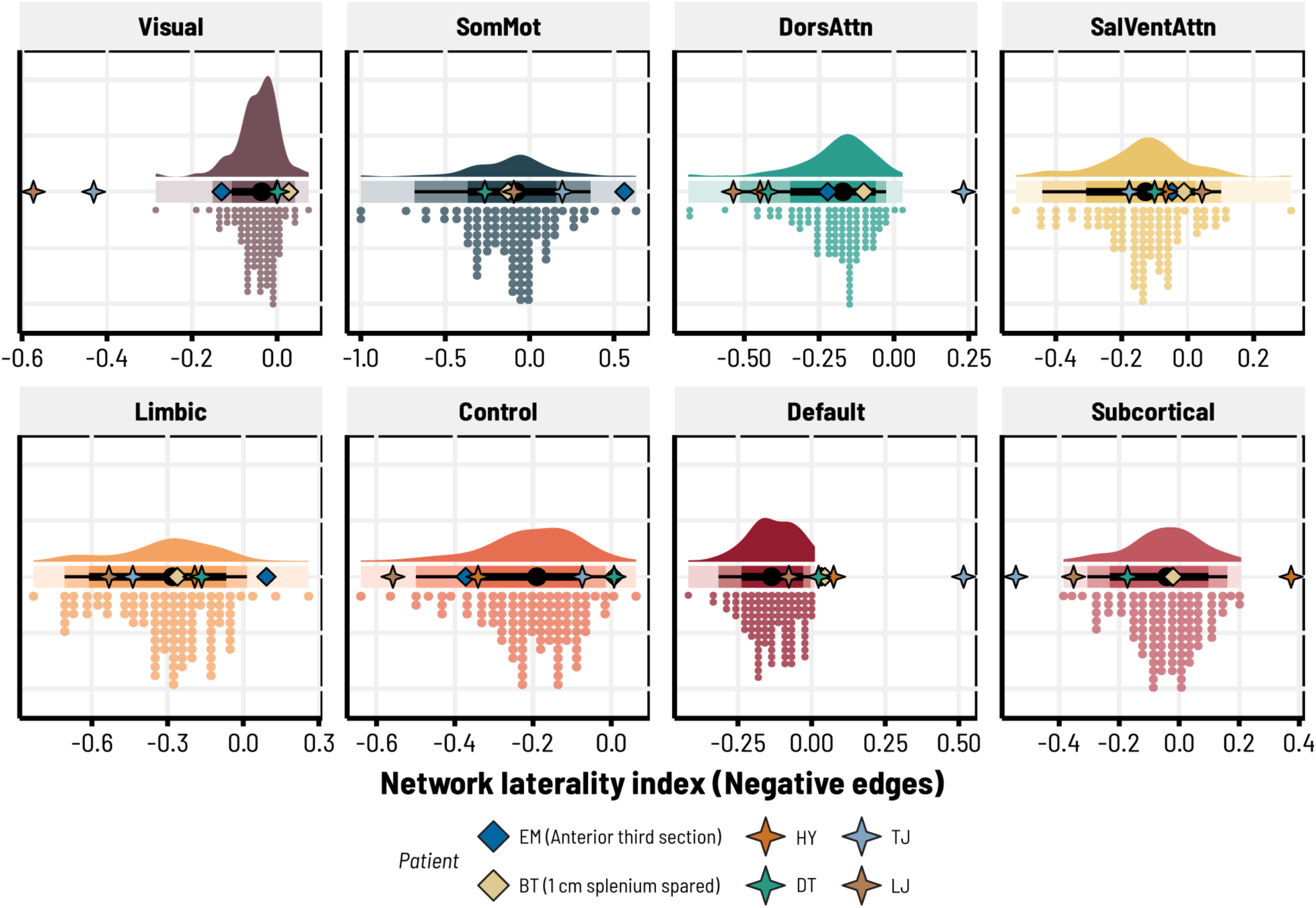
Anti-correlated functional connectivity is generally not more lateralized in callosotomy patients. Network laterality indices across seven canonical resting-state systems^17,18^ and subcortical regions^19^. To contrast against Fig. 3, here we estimate lateralization of negative (anti-correlated) connections. Values closer to 0 indicate an equal balance between intra- and inter-hemispheric (negative) FC, while values closer to -1 or 1 indicate more inter-hemispheric FC intra-hemispheric (negative) FC, respectively. Note that, unlike in Fig. 3, most distributions skew to the left (indicating that negative FC is predominantly inter-hemispheric, i.e., not lateralized), and both full and partial splits frequently fall within the range of HCP controls. Point intervals for each control distribution give the HCP100 median (black dot) and the 80-95% quantiles around the median (thick black lines and thin black lines, respectively). Abbreviations: FC; Functional Connectivity; SomMot, SomatoMotor Network; DorsAttn, Dorsal Attention Network; SalVentAttn, Salience/Ventral Attention Network.

**Extended Data Fig. 4.**
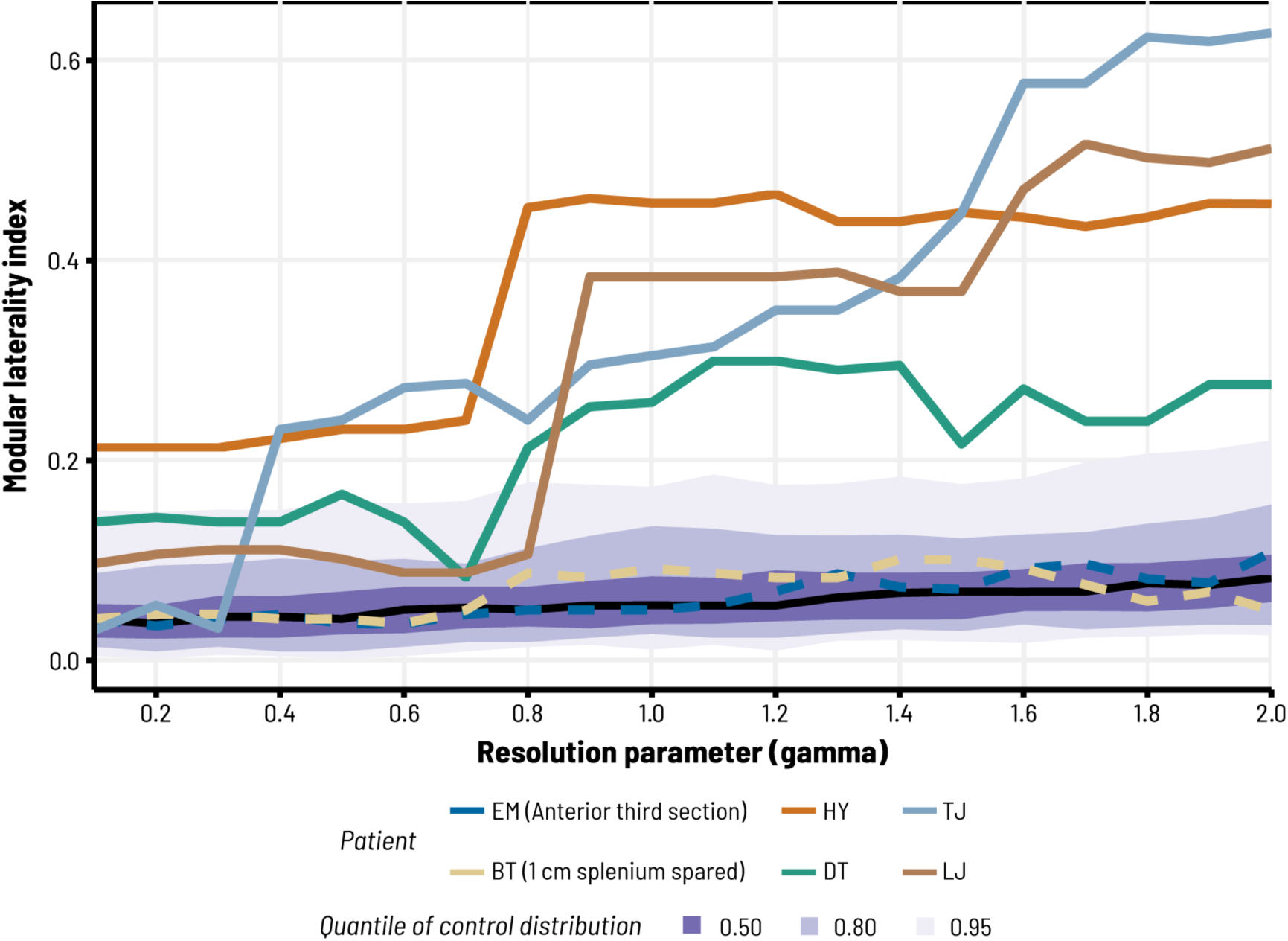
Lateralization of functional modules is readily-apparent in full splits regardless of resolution. Modularity maximization algorithms are sensitive to a spatial resolution parameter, γ^28^. A value of γ =1 is often treated as a ‘default’ level of resolution, whereas values of γ < 1 or γ > 1 produce progressively more coarse or fine-grained solutions, respectively. Here we show that differences in modular laterality between patients and healthy adult controls are not strictly-dependent upon the chosen value of γ. We performed 1000 iterations of modularity maximization at each of 20 levels of γ (ranging from [0.10, 2] in steps of 0.10), derived an optimal consensus partition at each level (see **Methods**), and estimated a laterality index for the resulting partitions^27^ (such that values closer to 1 indicate more lateralized network topologies). Full splits (solid-colored lines) rapidly diverged from the distribution of healthy controls (quantile intervals shown in gradations of purple, with the solid black line representing the median) demonstrating more lateralized organization of functional modules, even at coarse levels of γ < 1. Partial splits (dotted colored lines), by contrast, consistently remained within the control distribution.

**Extended Data Fig. 5.**
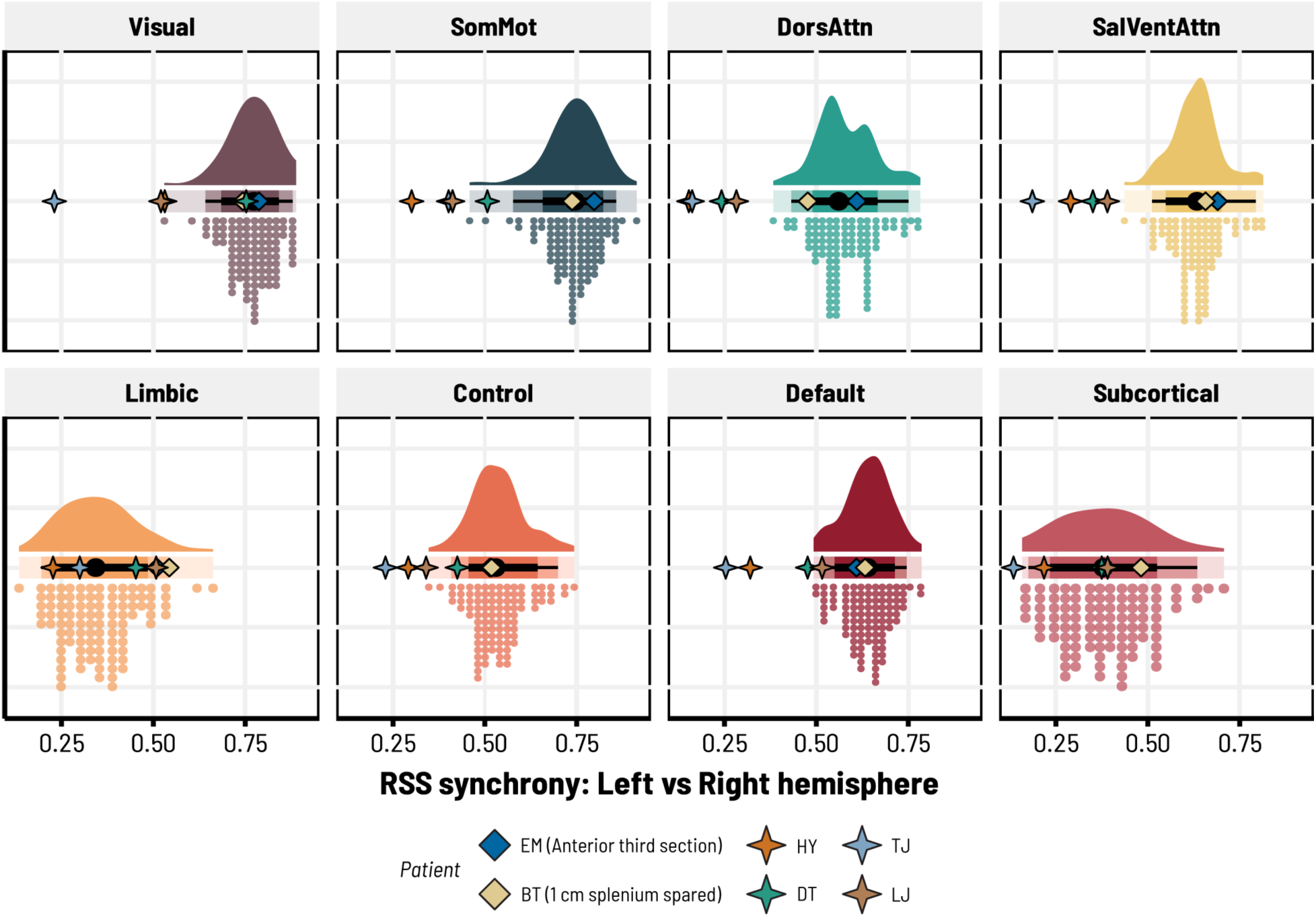
**Estimates of large-scale dynamic functional synchrony by network**. Decomposing whole-brain comparisons of RSS synchrony in Fig. 6b to individual networks reveals nuanced differences between patients and neurotypical adults (colored distributions). As before, partial splits consistently fall within the range of HCP controls. Full splits generally show more desynchronized network dynamics in higher-order cognitive systems, but comparatively-neurotypical levels of inter-hemispheric synchrony in limbic and subcortical networks (which are not strictly dependent upon structural connections through the corpus callosum, also reflecting Fig. 3). Point intervals under the control distributions give the HCP100 median (black dot) and 80-95% quantiles around the median (thick black lines and thin black lines, respectively). Abbreviations: RSS, Root-Sum-Of-Squares (see **Methods**); SomMot, SomatoMotor Network; DorsAttn, Dorsal Attention Network; SalVentAttn, Salience/Ventral Attention Network.

## Notes

### Competing Interest Statement

The authors have declared no competing interest.

### Summary of Updates

Manuscript reformatted for submission to Nature Neuroscience, with additional minor textual revisions from previous version

## References

1. Bassett DS, Sporns O. Network neuroscience. Nat Neurosci 20, 353–364 (2017).

2. Honey CJ, et al. Predicting human resting-state functional connectivity from structural connectivity. Proc Natl Acad Sci U S A 106, 2035–2040 (2009).

3. van den Heuvel MP, Mandl RC, Kahn RS, Hulshoff Pol HE. Functionally linked resting-state networks reflect the underlying structural connectivity architecture of the human brain. Hum Brain Mapp 30, 3127–3141 (2009).

4. Fukushima M, Betzel RF, He Y, van den Heuvel MP, Zuo XN, Sporns O. Structure-function relationships during segregated and integrated network states of human brain functional connectivity. Brain Struct Funct 223, 1091–1106 (2018).

5. Hermundstad AM, et al. Structurally-constrained relationships between cognitive states in the human brain. PLoS Comput Biol 10, e1003591 (2014).

6. Hermundstad AM, et al. Structural foundations of resting-state and task-based functional connectivity in the human brain. Proc Natl Acad Sci U S A 110, 6169–6174 (2013).

7. Hofer S, Frahm J. Topography of the human corpus callosum revisited--comprehensive fiber tractography using diffusion tensor magnetic resonance imaging. Neuroimage 32, 989–994 (2006).

8. Liu Y, et al. Connectivity-Based Topographical Changes of the Corpus Callosum During Aging. Front Aging Neurosci 13, 753236 (2021).

9. Hung SC, et al. Early recovery of interhemispheric functional connectivity after corpus callosotomy. Epilepsia 60, 1126–1136 (2019).

10. Johnston JM, et al. Loss of resting interhemispheric functional connectivity after complete section of the corpus callosum. J Neurosci 28, 6453–6458 (2008).

11. Roland JL, et al. On the role of the corpus callosum in interhemispheric functional connectivity in humans. Proc Natl Acad Sci U S A 114, 13278–13283 (2017).

12. Pizoli CE, et al. Resting-state activity in development and maintenance of normal brain function. Proc Natl Acad Sci U S A 108, 11638–11643 (2011).

13. Gazzaniga MS. Forty-five years of split-brain research and still going strong. Nat Rev Neurosci 6, 653–659 (2005).

14. Volz LJ, Gazzaniga MS. Interaction in isolation: 50 years of insights from split-brain research. Brain 140, 2051–2060 (2017).

15. Miller MB, Volz LJ, Simonson JM, Gazzaniga MS. Split-brain patients: A clinical versus experimental perspective. In: Cerebral Asymmetries (eds Papagno C, Corballis P). Elsevier BV (in press).

16. Bekir S, et al. No disconnection syndrome after near-complete callosotomy. Preprint at https://www.biorxiv.org/content/10.1101/2025.02.16.638524v1 (2025).

17. Schaefer A, et al. Local-Global Parcellation of the Human Cerebral Cortex from Intrinsic Functional Connectivity MRI. Cereb Cortex 28, 3095–3114 (2018).

18. Yeo BT, et al. The organization of the human cerebral cortex estimated by intrinsic functional connectivity. J Neurophysiol 106, 1125–1165 (2011).

19. Tian Y, Margulies DS, Breakspear M, Zalesky A. Topographic organization of the human subcortex unveiled with functional connectivity gradients. Nat Neurosci 23, 1421–1432 (2020).

20. Ciric R, et al. Benchmarking of participant-level confound regression strategies for the control of motion artifact in studies of functional connectivity. Neuroimage 154, 174–187 (2017).

21. Parkes L, Fulcher B, Yucel M, Fornito A. An evaluation of the efficacy, reliability, and sensitivity of motion correction strategies for resting-state functional MRI. Neuroimage 171, 415–436 (2018).

22. Catani M, Schotten M. Atlas of Human Brain Connections. Oxford University Press (2012).

23. Patel MD, Toussaint N, Charles-Edwards GD, Lin JP, Batchelor PG. Distribution and fibre field similarity mapping of the human anterior commissure fibres by diffusion tensor imaging. MAGMA 23, 399–408 (2010).

24. Di Virgilio G, Clarke S, Pizzolato G, Schaffner T. Cortical regions contributing to the anterior commissure in man. Exp Brain Res 124, 1–7 (1999).

25. Bassett DS, Gazzaniga MS. Understanding complexity in the human brain. Trends Cogn Sci 15, 200–209 (2011).

26. Sporns O, Betzel RF. Modular Brain Networks. Annu Rev Psychol 67, 613–640 (2016).

27. Doron KW, Bassett DS, Gazzaniga MS. Dynamic network structure of interhemispheric coordination. Proc Natl Acad Sci U S A 109, 18661–18668 (2012).

28. Fortunato S, Barthelemy M. Resolution limit in community detection. Proc Natl Acad Sci U S A 104, 36–41 (2007).

29. Pope M, Fukushima M, Betzel RF, Sporns O. Modular origins of high-amplitude cofluctuations in fine-scale functional connectivity dynamics. Proc Natl Acad Sci U S A 118, (2021).

30. Faskowitz J, Esfahlani FZ, Jo Y, Sporns O, Betzel RF. Edge-centric functional network representations of human cerebral cortex reveal overlapping system-level architecture. Nat Neurosci 23, 1644–1654 (2020).

31. Zamani Esfahlani F, et al. High-amplitude cofluctuations in cortical activity drive functional connectivity. Proc Natl Acad Sci U S A 117, 28393–28401 (2020).

32. Barttfeld P, Uhrig L, Sitt JD, Sigman M, Jarraya B, Dehaene S. Signature of consciousness in the dynamics of resting-state brain activity. Proc Natl Acad Sci U S A 112, 887–892 (2015).

33. Tooley UA, et al. The Age of Reason: Functional Brain Network Development during Childhood. J Neurosci 42, 8237–8251 (2022).

34. Sydnor VJ, et al. Neurodevelopment of the association cortices: Patterns, mechanisms, and implications for psychopathology. Neuron 109, 2820–2846 (2021).

35. Reynolds JE, Grohs MN, Dewey D, Lebel C. Global and regional white matter development in early childhood. Neuroimage 196, 49–58 (2019).

36. Whitaker KJ, et al. Adolescence is associated with genomically patterned consolidation of the hubs of the human brain connectome. Proc Natl Acad Sci U S A 113, 9105–9110 (2016).

37. Tyszka JM, Kennedy DP, Adolphs R, Paul LK. Intact bilateral resting-state networks in the absence of the corpus callosum. J Neurosci 31, 15154–15162 (2011).

38. Lassonde M, Sauerwein H, Chicoine AJ, Geoffroy G. Absence of disconnexion syndrome in callosal agenesis and early callosotomy: brain reorganization or lack of structural specificity during ontogeny? Neuropsychologia 29, 481–495 (1991).

39. Graham D, Tisdall MM, Gill D. Corpus callosotomy outcomes in pediatric patients: A systematic review. Epilepsia 57, 1053–1068 (2016).

40. Fuchs E, Flugge G. Adult neuroplasticity: more than 40 years of research. Neural Plast 2014, 541870 (2014).

41. Nomi JS, et al. Diffusion weighted imaging evidence of extra-callosal pathways for interhemispheric communication after complete commissurotomy. Brain Struct Funct 224, 1897–1909 (2019).

42. Uddin LQ, et al. Residual functional connectivity in the split-brain revealed with resting-state functional MRI. Neuroreport 19, 703–709 (2008).

43. Marcantoni I, et al. Interhemispheric functional connectivity: an fMRI study in callosotomized patients. Front Hum Neurosci 18, 1363098 (2024).

44. O’Reilly JX, et al. Causal effect of disconnection lesions on interhemispheric functional connectivity in rhesus monkeys. Proc Natl Acad Sci U S A 110, 13982–13987 (2013).

45. Funnell MG, Corballis PM, Gazzaniga MS. Cortical and subcortical interhemispheric interactions following partial and complete callosotomy. Arch Neurol 57, 185–189 (2000).

46. Gazzaniga MS, Freedman H. Observations on visual processes after posterior callosal section. Neurology 23, 1126–1130 (1973).

47. Risse GL, Gates J, Lund G, Maxwell R, Rubens A. Interhemispheric transfer in patients with incomplete section of the corpus callosum. Anatomic verification with magnetic resonance imaging. Arch Neurol 46, 437–443 (1989).

48 Gazzaniga MS. Interhemispheric integration. In: Neurobiology of Neocortex (eds Rakic P, Singer W). John Wiley & Sons (1988).

49. Seguin C, Mansour LS, Sporns O, Zalesky A, Calamante F. Network communication models narrow the gap between the modular organization of structural and functional brain networks. Neuroimage 257, 119323 (2022).

50. Suarez LE, Markello RD, Betzel RF, Misic B. Linking Structure and Function in Macroscale Brain Networks. Trends Cogn Sci 24, 302–315 (2020).

51. Misic B, et al. Network-Level Structure-Function Relationships in Human Neocortex. Cereb Cortex 26, 3285–3296 (2016).

52. Aboitiz F, Lopez J, Montiel J. Long distance communication in the human brain: timing constraints for inter-hemispheric synchrony and the origin of brain lateralization. Biol Res 36, 89–99 (2003).

53. Aboitiz F, Scheibel AB, Fisher RS, Zaidel E. Fiber composition of the human corpus callosum. Brain Res 598, 143–153 (1992).

54. Horowitz A, Barazany D, Tavor I, Bernstein M, Yovel G, Assaf Y. In vivo correlation between axon diameter and conduction velocity in the human brain. Brain Struct Funct 220, 1777–1788 (2015).

55. van den Heuvel MP, Sporns O. Rich-club organization of the human connectome. J Neurosci 31, 15775–15786 (2011).

56. Hagmann P, et al. Mapping the structural core of human cerebral cortex. PLoS Biol 6, e159 (2008).

57. Koch C, Massimini M, Boly M, Tononi G. Neural correlates of consciousness: progress and problems. Nat Rev Neurosci 17, 307–321 (2016).

58. Melloni L, et al. An adversarial collaboration protocol for testing contrasting predictions of global neuronal workspace and integrated information theory. PLoS One 18, e0268577 (2023).

59. Koch C, Massimini M, Boly M, Tononi G. Posterior and anterior cortex - where is the difference that makes the difference? Nat Rev Neurosci 17, 666 (2016).

60. Van Essen DC, et al. The WU-Minn Human Connectome Project: an overview. Neuroimage 80, 62–79 (2013).

61. Van Essen DC, et al. The Human Connectome Project: a data acquisition perspective. Neuroimage 62, 2222–2231 (2012).

62. Tustison NJ, et al. The ANTsX ecosystem for quantitative biological and medical imaging. Sci Rep 11, 9068 (2021).

63. Tustison NJ, et al. Large-scale evaluation of ANTs and FreeSurfer cortical thickness measurements. Neuroimage 99, 166–179 (2014).

64. Evans AC, Janke AL, Collins DL, Baillet S. Brain templates and atlases. Neuroimage 62, 911–922 (2012).

65. Jenkinson M, Beckmann CF, Behrens TE, Woolrich MW, Smith SM. Fsl. *Neuroimage* **62**, 782–790 (2012).

66. Hua K, et al. Tract probability maps in stereotaxic spaces: analyses of white matter anatomy and tract-specific quantification. Neuroimage 39, 336–347 (2008).

67. Wakana S, et al. Reproducibility of quantitative tractography methods applied to cerebral white matter. Neuroimage 36, 630–644 (2007).

68. Friston KJ, Rotshtein P, Geng JJ, Sterzer P, Henson RN. A critique of functional localisers. Neuroimage 30, 1077–1087 (2006).

69. Zamani Esfahlani F, et al. Modularity maximization as a flexible and generic framework for brain network exploratory analysis. Neuroimage 244, 118607 (2021).

70. Rubinov M, Sporns O. Weight-conserving characterization of complex functional brain networks. Neuroimage 56, 2068–2079 (2011).

71. Rubinov M, Sporns O. Complex network measures of brain connectivity: uses and interpretations. Neuroimage 52, 1059–1069 (2010).

